# Species-specific susceptibility and low transmission of the historical Japanese encephalitis virus Nakayama strain in North American *Culex* mosquitoes

**DOI:** 10.64898/2026.08.10.744046

**Authors:** RL Fay, EM Banker, AF Payne, AP Dupuis, J Stout, A Russell, V Schnurr, SM Bialosuknia, L Munn, EA Mordecai, AT Ciota

## Abstract

Japanese encephalitis virus (JEV) is an emerging mosquito-borne flavivirus with potential for geographic expansion, yet the risk of establishment in North America remains poorly characterized. We assessed vector competence of three North American *Culex* species (*Cx. pipiens, Cx. quinquefasciatus, and Cx. tarsalis*) for the JEV Nakayama strain, isolated from human brain in 1934 in Japan, across five constant temperatures (15, 20, 25, 30, and 33°C) at 4, 7, and 14 days post-feeding, quantifying infection, dissemination, and transmission rates. Vector competence was low but non-zero across all species. *Cx. pipiens* showed higher infection rates than the other species, whereas *Cx. quinquefasciatus* and *Cx. tarsalis* were minimally susceptible under these experimental conditions. Temperature had limited effects on infection and no detectable effects on dissemination or transmission. These findings suggest limited transmission potential of JEV Nakayama in North America, with *Cx. pipiens* as a relatively permissive vector.

## Introduction

Japanese encephalitis virus (JEV) belongs to the genus *Orthoflavivirus*, a group of positive-sense, single-stranded RNA viruses within the family *Flaviviridae*. First isolated in Japan in 1935, JEV occurs in an enzootic transmission cycle between waterbirds and Culex spp. mosquitoes. Epizootic transmission involves swine as amplifying hosts and humans and horses as incidental hosts^1,2^. An estimated 68,000 JEV cases occur annually^3^. Approximately 1% of humans develop febrile illness, with a mortality rate of 30% among these individuals2. JEV is vaccine-preventable: researchers developed an inactivated vaccine in the 1930s, like the vaccine administered today, which is ~90% effective^4^. Vaccination is limited to JEV-endemic regions, leaving populations in introduced areas highly susceptible. JEV is classified into five genotypes (GI–GV), with GI and GIII responsible for most contemporary human infections^1^. JEV is predominantly found in subtropical regions including Asia and the Western Pacific^5^. Despite immunization campaigns, the virus continues to circulate in southern Asia^6^. This raises concerns for JEV risk in regions with competent vectors and hosts, as well as geographic expansion under environmental change^7^.

In the last three decades, zoonotic diseases have emerged in new regions, a process linked to globalization and climate change. Vector-borne diseases pose a threat as they are sensitive to climate and can cause explosive epidemics. Emergence events have been linked to favorable environmental conditions and competent vectors, including the West Nile virus (WNV) introduction to the United States (US) in 1999, and chikungunya and Zika virus outbreaks in the Americas in the 2000s and 2015, respectively^8–11^. In 2022, local transmission of JEV was identified for the first time in Australia, demonstrating its potential for JEV expansion into new regions^12^. During this outbreak, 45 human cases and seven deaths were reported, while reproductive losses in pig herds imposed economic costs on the swine industry^12^. The presence of competent vectors and hosts puts the US at risk for JEV introduction^13–15^. Early detection may be hindered in the US by limited surveillance testing in vectors, wildlife, domestic animals, and humans, and flavivirus cross-reactivity, which can make novel virus circulation difficult to detect.

Vector-borne disease transmission is highly sensitive to temperature, with transmission generally occurring within species-specific thermal limits defined by minimum, optimum, and maximum temperatures for pathogen transmission^16^. However, only limited work has examined the effects of temperature on JEV transmission^17^, and none has focused on North American mosquito vectors. Here, we experimentally evaluate the vector competence of three North American Culex mosquito species across a range of temperatures using the well-characterized Nakayama genotype GIII reference strain of JEV to assess the thermal conditions under which transmission may be possible. By characterizing temperature-dependent variation in JEV vector competence, this study evaluates the potential for JEV transmission in the US and provides information to guide public health preparedness and risk assessment following potential introduction.

## Methods

This study used the Nakayama (EF571853) JEV strain, originally isolated from human brain tissue in Tokyo, Japan, in 1934 (P9, Vero 4), a Genotype III strain. *Cx. pipiens* (Suffolk County, NY), *Cx. quinquefasciatus* (Horry County, SC), and *Cx. tarsalis* (Coachella Valley, CA) colonies were maintained in 30.5-cm^3^ cages at 27 ± 2°C, 45–65% relative humidity, and a 16:8-h light:dark cycle, with 10% sucrose provided *ad libitum*.

For vector competence assays, 4–7-day-old females were fed JEV blood meals containing 5.9–7.3 log_10_PFU/mL (*Cx. tarsalis, 7.3; Cx. quinquefasciatus, 6.1; Cx. pipiens*, 5.9 log_10_PFU/mL). Blood meals consisted of a 4:1 mixture of diluted virus and defibrinated chicken blood (Rockland Immunochemicals, Inc., Pottstown, PA) with 2.5% sucrose. Mosquitoes were fed for 1 h at 37°C using an artificial feeding system (Hemotek, Blackburn, UK), anesthetized with CO_2_, and engorged females were held at 15, 20, 25, 30, or 33°C. At 4, 7, and 12 days post-feeding (dpf), 26–40 mosquitoes per species-temperature combination were assayed for infection, dissemination, and transmission by detecting virus in bodies, legs, and saliva, respectively. Saliva was collected for 30 min using capillary tubes containing ~20 µL fetal bovine serum and 50% sucrose and transferred to 125 µL mosquito diluent (PBS, 20% FBS, 50 µg/mL penicillin/streptomycin, 50 µg/mL gentamicin, and 2.5 µg/mL amphotericin B). Bodies and legs were stored separately in 500 µL mosquito diluent with a 4.5-mm zinc-plated steel ball (Daisy, Dallas, TX) at −80°C. Experiments were conducted in three independent groups, with each group representing one mosquito species.

Body and leg samples were homogenized separately at 30 Hz for 30 s and centrifuged at 12,000 rpm for 3 min at 4°C. RNA was extracted using the MagMAX-96 Viral RNA Isolation Kit and KingFisher Flex 96 instrument (Thermo Fisher Scientific, Waltham, MA). Viral RNA was quantified by one-step RT-qPCR using qScript XLT One-Step RT-qPCR ToughMix, Low ROX (QuantaBio, Beverly, MA) on a QuantStudio 5 Real-Time PCR System (Thermo Fisher Scientific). Universal primers described by Shao et al. (2018)18 were used, with PFU/mL standards for quantification. Mosquitoes with visible residual blood meals on day 4 were excluded from infection analyses. Data were analyzed using GraphPad Prism 11 and RStudio.

JEV blood meal titers were determined by plaque assay. Confluent Vero cell monolayers in six-well plates were inoculated with 100 µL of 10-fold serial dilutions of virus for 1 h at 37°C. Cells were overlaid with 3 mL of a 1:1 mixture of 2× EMEM containing 10% FBS and 1.2% Oxoid agar and incubated at 37°C for 4 days. Plates were then overlaid with 1:1 2× EMEM containing 2% FBS and agar with neutral red (1.5% final concentration). Plaques were counted 24 h later.

## Results

To assess the potential for JEV transmission in the US, we evaluated the vector competence of three North American Culex species: *Cx. tarsalis, Cx. quinquefasciatus*, and *Cx. pipiens*. Adult mosquitoes were fed an infectious blood meal containing the historical Nakayama strain of JEV and maintained at 15, 20, 25, 30, or 33°C. Infection, dissemination, and transmission were assessed at 4, 7, and 14 dpf.

Overall, all three species exhibited low vector competence across temperatures. Infection prevalence never exceeded 37.5%, while dissemination and transmission remained below 10% and 2.5%, respectively, among exposed mosquitoes across temperature treatments (Fig. 1; Table 1). When data were pooled across sampling time points, logistic regression identified significant overall effects of mosquito species (LR χ^2^ = 28.26, df = 2, *P* < 0.001) and temperature (LR χ^2^ = 15.08, df = 4, *P* = 0.0045) on infection prevalence, whereas the species × temperature interaction was not significant (LR χ^2^ = 13.92, df = 8, *P* = 0.084). Post hoc comparisons indicated that *Cx. pipiens* had significantly higher infection prevalence than *Cx. quinquefasciatus* at 15, 20, 25, and 30°C and than *Cx. tarsalis* at 15, 20, and 30°C (Fig. 1A). No significant differences among species were detected at 33°C. In contrast, logistic regression detected no significant effects of species, temperature, or their interaction on dissemination or transmission prevalence (all *P* > 0.10), consistent with low dissemination and transmission rates observed across all species and temperature treatments (Fig. 1B–C).

**Table 1.** Vector competence results of North American *Culex* mosquitoes for JEV across different temperatures. Percentages are calculated from the total number of mosquitoes tested. Columns are grouped by assay day post-feeding (dpf = 4, 7, or 14); rows are grouped by mosquito species. For infection, dissemination, and transmission, the number of positive individuals is given in the first column and the percentage in the second column. Only mosquitoes that were infected were tested for dissemination, and only those with dissemination were tested for transmission.

| *Cx. pipiens* | D4 |  |  |  |  |  | D7 |  |  |  |  |  | D14 |  |  |  |  |  |
| --- | --- | --- | --- | --- | --- | --- | --- | --- | --- | --- | --- | --- | --- | --- | --- | --- | --- | --- |
| Temperature | Infection | % | dissemination | % | Transmission | % | Infection | % | dissemination | % | Transmission | % | Infection | % | dissemination | % | Transmission | % |
| 15 | 11 | 27.50% | 1 | 2.50% | 1 | 2.50% | 14 | 35.00% | 1 | 2.50% | 1 | 2.50% | 10 | 25.00% | 1 | 2.50% | 0 | 0.00% |
| 20 | 15 | 37.50% | 4 | 10.00% | 1 | 2.50% | 14 | 35.00% | 1 | 2.50% | 0 | 0.00% | 10 | 25.00% | 0 | 0.00% | 0 | 0.00% |
| 25 | 14 | 35.00% | 2 | 5.00% | 0 | 0.00% | 6 | 15.00% | 2 | 5.00% | 0 | 0.00% | 10 | 25.00% | 3 | 7.50% | 1 | 2.50% |
| 30 | 3 | 7.50% | 1 | 2.50% | 1 | 2.50% | 14 | 35.00% | 2 | 5.00% | 2 | 5.00% | 5 | 16.67% | 0 | 0.00% | 0 | 0.00% |
| 33 | 7 | 17.50% | 1 | 2.50% | 1 | 2.50% | 3 | 7.50% | 2 | 5.00% | 1 | 2.50% | N/A | N/A | N/A | N/A | N/A | N/A |
| *Cx. quinquefasciatus* | D4 |  |  |  |  |  | D7 |  |  |  |  |  | D14 |  |  |  |  |  |
| Temperature | Infection | % | dissemination | % | Transmission | % | Infection | % | dissemination | % | Transmission | % | Infection | % | dissemination | % | Transmission | % |
| 15 | 0 | 0.00% | 0 | 0.00% | 0 | 0.00% | 4 | 10.00% | 0 | 0.00% | 0 | 0.00% | 2 | 5.00% | 0 | 0.00% | 0 | 0.00% |
| 20 | 5 | 12.50% | 0 | 0.00% | 0 | 0.00% | 9 | 22.50% | 0 | 0.00% | 0 | 0.00% | 4 | 10.00% | 0 | 0.00% | 0 | 0.00% |
| 25 | 6 | 15.00% | 1 | 2.50% | 0 | 0.00% | 6 | 15.00% | 1 | 2.50% | 0 | 0.00% | 2 | 5.00% | 0 | 0.00% | 0 | 0.00% |
| 30 | 2 | 5.00% | 0 | 0.00% | 0 | 0.00% | 2 | 5.00% | 0 | 0.00% | 0 | 0.00% | 0 | 0.00% | 0 | 0.00% | 0 | 0.00% |
| 33 | 1 | 2.50% | 1 | 2.50% | 0 | 0.00% | 3 | 7.50% | 0 | 0.00% | 0 | 0.00% | 0 | 0.00% | 0 | 0.00% | 0 | 0.00% |
| *Cx. tarsalis* | D4 |  |  |  |  |  | D7 |  |  |  |  |  | D14 |  |  |  |  |  |
| Temperature | Infection | % | dissemination | % | Transmission | % | Infection | % | dissemination | % | Transmission | % | Infection | % | dissemination | % | Transmission | % |
| 15 | 10 | 25.00% | 0 | 0.00% | 0 | 0.00% | 4 | 10.00% | 1 | 2.50% | 0 | 0.00% | 1 | 2.50% | 0 | 0.00% | 0 | 0.00% |
| 20 | 6 | 15.00% | 1 | 2.50% | 0 | 0.00% | 2 | 5.00% | 1 | 2.50% | 0 | 0.00% | 2 | 5.00% | 1 | 2.50% | 0 | 0.00% |
| 25 | 8 | 20.00% | 1 | 2.50% | 0 | 0.00% | 9 | 22.50% | 3 | 7.50% | 1 | 2.50% | 3 | 7.50% | 1 | 2.50% | 1 | 2.50% |
| 30 | 2 | 5.00% | 0 | 0.00% | 0 | 0.00% | 4 | 10.00% | 1 | 2.50% | 0 | 0.00% | 1 | 2.50% | 0 | 0.00% | 0 | 0.00% |
| 33 | 2 | 5.00% | 1 | 2.50% | 1 | 2.50% | 1 | 2.50% | 0 | 0.00% | 0 | 0.00% | 5 | 19.23% | 1 | 2.50% | 0 | 0.00% |

**Figure 1.**
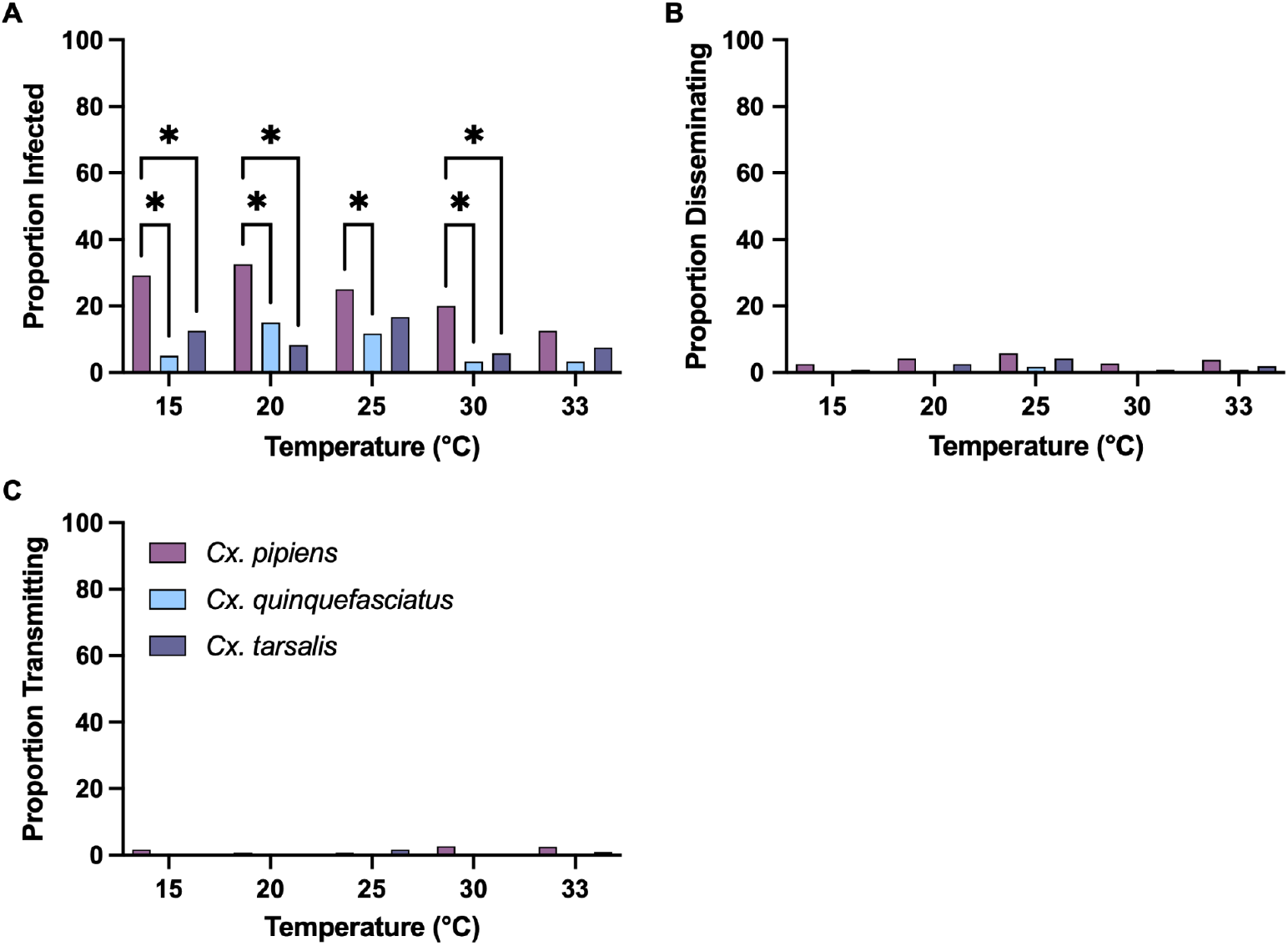
Vector competence of three North American *Culex* mosquito species for JEV across temperatures. (**A–C**) Proportion of mosquitoes infected, with disseminated infection, and transmitting JEV, respectively, pooled across 4, 7, and 14 dpf. *Cx. pipiens* is shown in pink, *Cx. quinquefasciatus* in blue, and *Cx. tarsalis* in purple. Brackets indicate significant pairwise differences among mosquito species within each temperature based on Tukey-adjusted post hoc comparisons from the logistic regression model (*P* < 0.05). Logistic regression revealed significant overall effects of species (LR χ^2^ = 28.26, df = 2, *P* < 0.001) and temperature (LR χ^2^ = 15.08, df = 4, *P* = 0.0045) on infection prevalence, whereas the species × temperature interaction was not significant (LR χ^2^ = 13.92, df = 8, *P* = 0.084). Neither species, temperature, nor their interaction significantly affected dissemination or transmission (all *P* > 0.10).

We next examined viral titers within infected mosquitoes across temperatures (Fig. 2). *Cx. quinquefasciatus* and *Cx. pipiens* exhibited significantly higher viral loads at 33°C than at all other temperatures, except *Cx. quinquefasciatus* at 30°C, which did not differ significantly from 33°C (Fig. 2BC). In contrast, viral loads in *Cx. tarsalis* did not differ significantly across temperatures (Fig. 2A).

**Figure 2.**
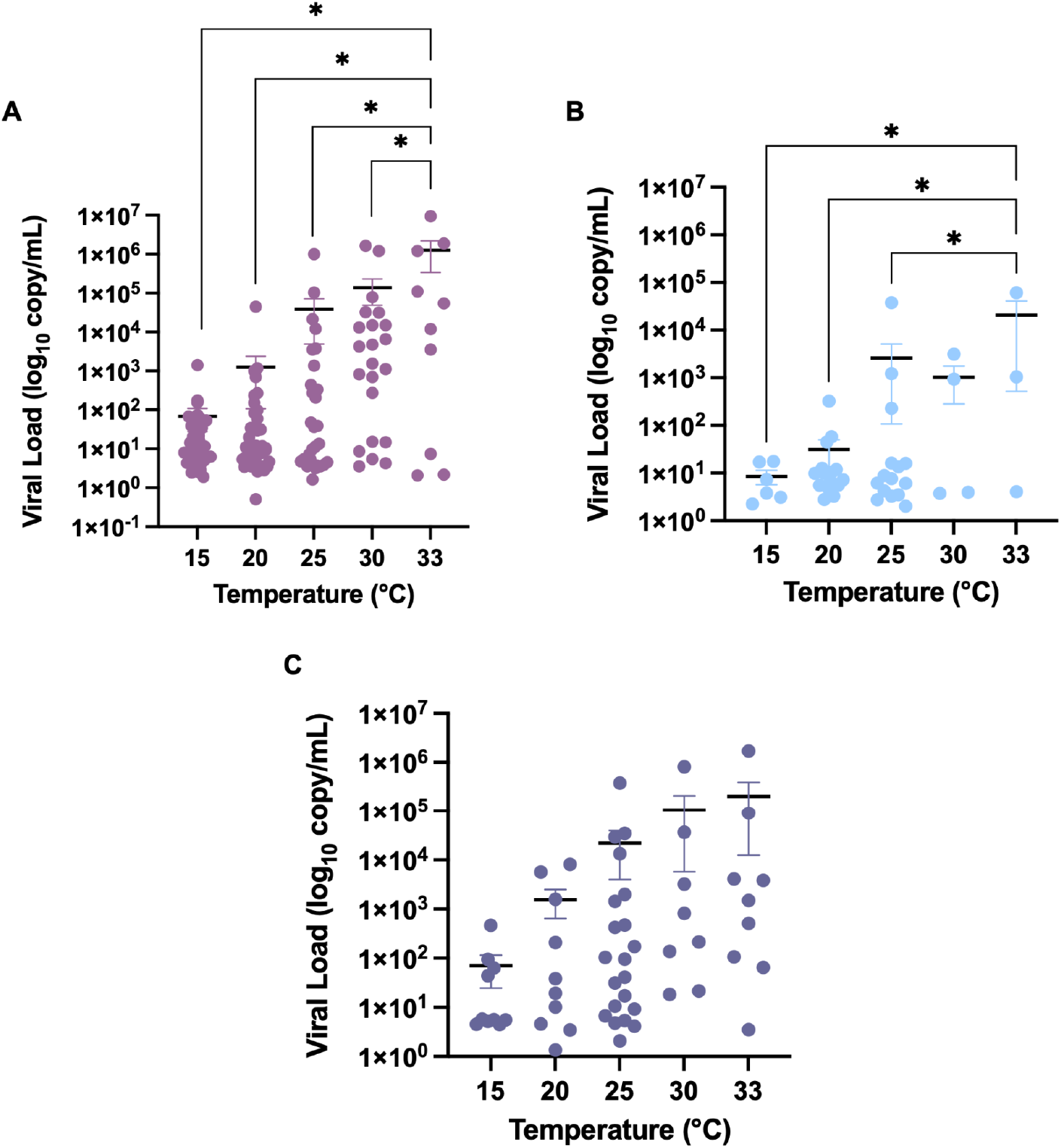
JEV viral load in North American *Culex* mosquitoes. (**A–C**) JEV viral load in the bodies of infected *Cx. pipiens, Cx. quinquefasciatus*, and *Cx. tarsalis*, respectively. JEV viral load (genome copy number equivalents) was quantified in individual mosquito bodies using JEV-specific quantitative reverse transcription PCR and a standard curve. Viral loads at 4, 7, and 14 dpf are combined with data points shown for individual mosquitoes, with bars indicating the mean ± SEM. Statistical significance was determined using one-way ANOVA followed by Tukey’s multiple comparisons test (*p<0.05).

## Discussion

Our findings suggest that the historical Nakayama strain of JEV has limited potential for transmission by the three North American *Culex* species examined, indicating a low likelihood of establishment following introduction into the US. Infection rates were low across temperatures, with *Cx. pipiens* exhibiting the greatest susceptibility and significantly higher infection rates than *Cx. quinquefasciatus* at all temperatures except 33°C, and than *Cx. tarsalis* at 15, 20, and 30°C (Fig. 1 and Table 1). Infection differed significantly among species overall, whereas temperature and the species-by-temperature interaction had little effect on infection. Dissemination and transmission remained rare with no significant effects of species or temperature on these outcomes. Despite low infection, dissemination, and transmission rates, viral titers in infected *Cx. quinquefasciatus* and *Cx. pipiens* increased with temperature, with significantly higher viral loads at 33°C than at lower temperatures (Fig. 2). This pattern is consistent with studies of other flaviviruses, where higher temperatures accelerate viral replication in mosquitoes^19^.

Although this study evaluated only a single JEV strain, vector competence may differ among viral genotypes. The Nakayama strain belongs to genotype III, whereas genotype IV has recently been implicated in outbreaks and geographic expansion^20^. Mosquito populations were laboratory-colonized, and vector competence may differ among field populations because of genetic and environmental variation^21^. Future studies should therefore evaluate contemporary JEV genotypes and field-derived North American mosquito populations across temperatures.

To our knowledge, this is the first study to assess temperature effects on JEV vector competence in North American mosquito vectors. These data provide an important foundation for defining vector competence and thermal suitability of JEV transmission in the US. Incorporating these estimates into mechanistic transmission models will improve predictions of where and when JEV establishment may be possible.

## Acknowledgments

We thank the NYS Arbovirology Laboratory insectary staff for support and assistance. We thank the Wadsworth Center Media and Tissue Culture Facility for providing cells and media. RLF was funded by the National Institutes of Health 1F32AI191534-01 and R35GM133439. ATC was supported by National Institutes of Health R01AI168097. EAM was supported by grants from the National Science Foundation (DEB-2011147 with Fogarty International Center) and the National Institutes of Health (R35GM133439, R01AI168097).

## Supplemental Figure

**Figure S1.**
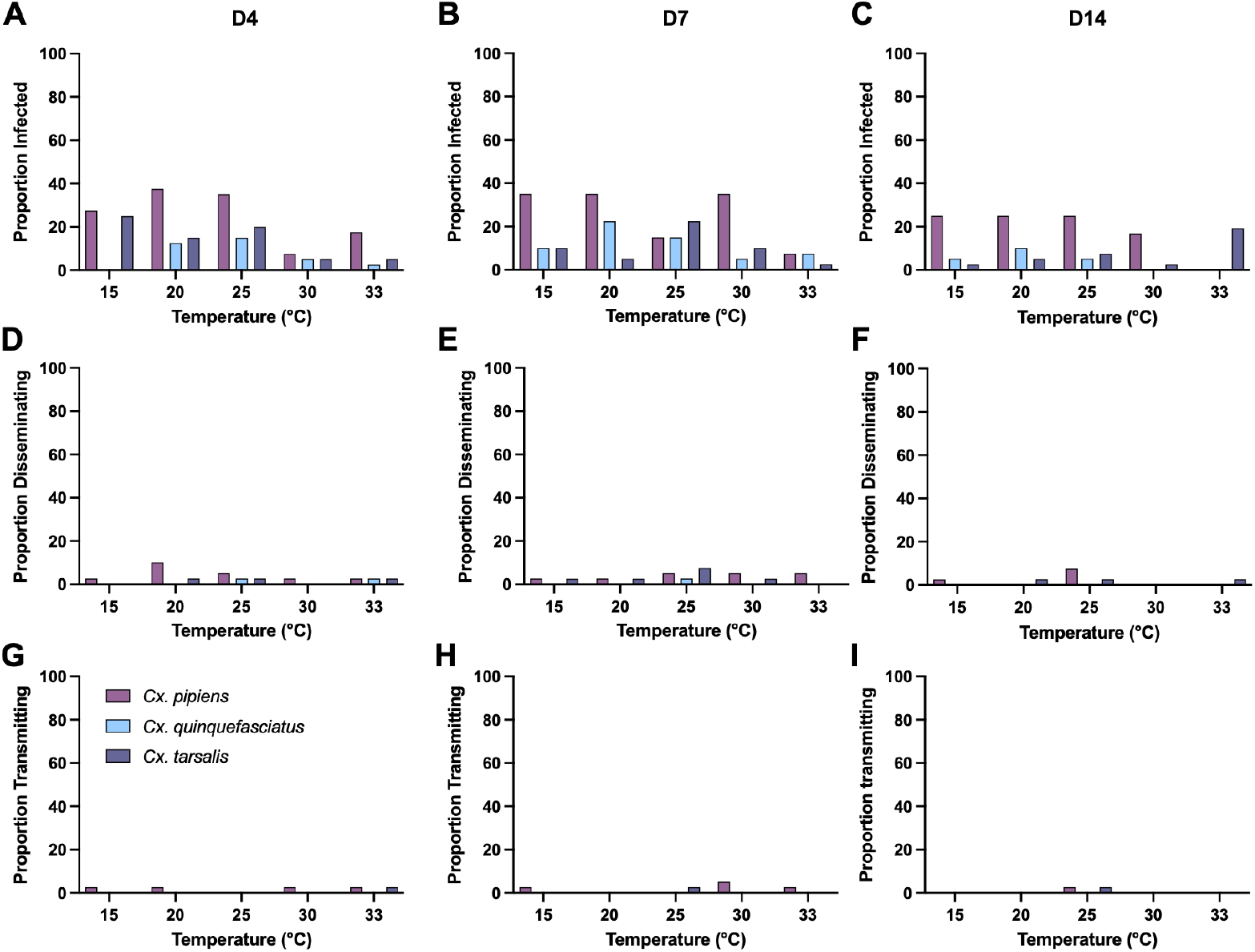
Time-point-specific vector competence of three North American *Culex* mosquito species for JEV across temperature. (**A–C**) Proportion of mosquitoes infected with JEV at 4, 7, and 14 dpf, respectively. (**D–F**) Proportion of mosquitoes with disseminated JEV infections at 4, 7, and 14 dpf, respectively. (**G–I**) Proportion of mosquitoes transmitting JEV at 4, 7, and 14 dpf, respectively. *Cx. pipiens* shown in pink, *Cx. quinquefasciatus* in blue, and *Cx. tarsalis* in purple.

## Notes

### Competing Interest Statement

The authors have declared no competing interest.

## References

1. Solomon T, Ni H, Beasley DWC, Ekkelenkamp M, Cardosa MJ, Barrett ADT., 2003. Origin and evolution of Japanese encephalitis virus in Southeast Asia. J Virol 77: 3091–3098

2. Halstead SB, Jacobson J., 2003. Japanese encephalitis. Adv Virus Res 61: 103–138

3. Campbell G, Hills S, Fischer M, Jacobson J, Hoke C, Hombach J, Marfin A, Solomon T, Tsai T, Tsui V, Ginsburg A., 2011. Estimated global incidence of Japanese encephalitis: Bull World Heal Organ 89: 766–774

4. Paulke-Korinek M, Kollaritsch H., 2008. Japanese encephalitis and vaccines: past and future prospects. Wien Klin Wochenschr 120: 15–19

5. Mackenzie JS, Williams DT, Smith DW., 2006. Japanese Encephalitis Virus: The geographic distribution, incidence, and spread of a virus with a propensity to emerge in new areas. Perspect Méd Virol 16: 201–268

6. (CDC) C for DC and P., 2013. Japanese encephalitis surveillance and immunization--Asia and the Western Pacific, 2012. MMWR Morb Mortal Wkly Rep 62: 658–62

7. Skinner EB, Sartorius B, Furuya-Kanamori L, Craig AT, Kiani B, Johnson BJ, Moore KT, Hickson RI, Mordecai EA, Devine G, Lau CL., 2025. Ecological suitability of Japanese encephalitis virus in Australia: A modelling analysis of vector-host transmission dynamics to potential spillover in humans. PLOS Neglected Trop Dis 19: e0013722

8. Kramer LD, Ciota AT, Kilpatrick AM., 2019. Introduction, spread, and establishment of West Nile virus in the Americas. J Med Entomol 56: 1448–1455

9. Yactayo S, Staples JE, Millot V, Cibrelus L, Ramon-Pardo P., 2016. Epidemiology of Chikungunya in the Americas. J Infect Dis 214: S441–S445

10. Caminade C, Turner J, Metelmann S, Hesson JC, Blagrove MSC, Solomon T, Morse AP, Baylis M., 2017. Global risk model for vector-borne transmission of Zika virus reveals the role of El Niño 2015. Proc Natl Acad Sci 114: 119–124

11. Metsky HC, Matranga CB, Wohl S, Schaffner SF, Freije CA, Winnicki SM, West K, Qu J, Baniecki ML, Gladden-Young A, Lin AE, Tomkins-Tinch CH, Ye SH, Park DJ, Luo CY, et al., 2017. Zika virus evolution and spread in the Americas. Nature 546: 411–415

12. Hurk AF van den, Skinner E, Ritchie SA, Mackenzie JS., 2022. The emergence of Japanese encephalitis virus in Australia in 2022: Existing Knowledge of Mosquito Vectors. Viruses 14: 1208

13. Kramer LD, Chin P, Cane RP, Kauffman EB, Mackereth G., 2011. Vector competence of New Zealand mosquitoes for selected arboviruses. Am J Trop Med Hyg 85: 182–189

14. Reeves WC, Hammon WM, Espana W the TA of GGW and C., 1946. Laboratory transmission of Japanese B encephalitis virus by seven species (three genera) of North American mosquitoes. J Exp Med 83: 185–194

15. Huang Y-JS, Harbin JN, Hettenbach SM, Maki E, Cohnstaedt LW, Barrett ADT, Higgs S, Vanlandingham DL., 2015. Susceptibility of a North American Culex quinquefasciatus to Japanese encephalitis Virus. Vector-Borne Zoonotic Dis 15: 709–711

16. Mordecai EA, Caldwell JM, Grossman MK, Lippi CA, Johnson LR, Neira M, Rohr JR, Ryan SJ, Savage V, Shocket MS, Sippy R, Ibarra AMS, Thomas MB, Villena O., 2019. Thermal biology of mosquito-borne disease. Ecol Lett 22: 1690–1708

17. Folly AJ, Dorey-Robinson D, Hernández-Triana LM, Ackroyd S, Vidana B, Lean FZX, Hicks D, Nuñez A, Johnson N., 2021. Temperate conditions restrict Japanese encephalitis virus infection to the mid-gut and prevents systemic dissemination in Culex pipiens mosquitoes. Sci Rep-uk 11: 6133

18. Shao N, Li F, Nie K, Fu SH, Zhang WJ, He Y, Lei WW, Wang QY, Liang GD, Cao YX, Wang HY., 2018. TaqMan real-time RT-PCR assay for detecting and differentiating Japanese encephalitis virus. Biomed Environ Sci 31: 208–214

19. Fay RL, Cruz-Loya M, Maffei JG, Mordecai EA, Ciota AT., 2025. Rising temperatures contribute to West Nile virus diversification and increased transmission potential. Sci Rep 15: 25016

20. Howard-Jones AR, Pham D, Jeoffreys N, Eden J-S, Hueston L, Kesson AM, Nagendra V, Samarasekara H, Newton P, Chen SC-A, O’Sullivan MV, Maddocks S, Dwyer DE, Kok J., 2022. Emerging genotype IV Japanese encephalitis virus outbreak in New South Wales, Australia. Viruses 14: 1853

21. Fay RL, Cruz-Loya M, Keyel AC, Price DC, Zink SD, Mordecai EA, Ciota AT., 2024. Population-specific thermal responses contribute to regional variability in arbovirus transmission with changing climates. iScience: 109934

